# StoPred: Accurate Stoichiometry Prediction for Protein Complexes Using Protein Language Models and Graph Attention

**DOI:** 10.1101/2025.10.20.683515

**Authors:** Quancheng Liu, Chunxiang Peng, Wei Zheng, Chengxin Zhang, Lydia Freddolino

## Abstract

Proteins often function as part of complexes, and the specific stoichiometry of these assemblies is critical for their biological roles, but experimental determination of assembly composition remains challenging and existing computational methods for stoichiometry prediction are limited. Existing approaches rely on template-based searches or require predefined stoichiometry for structure prediction, hampering their applicability to proteins without close homologs or known assembly states. Recent advances using protein language models (pLM) have enabled sequence-based prediction of homo-oligomer stoichiometry, but these methods are not applicable to hetero-oligomeric complexes and do not fully leverage inter-subunit relationships. Here, we present StoPred, a method that predicts the stoichiometry of protein complexes by integrating pLM embeddings with a graph attention network to model subunit-level interactions. StoPred infers stoichiometry directly from sequence or structure features for both homo-and hetero-oligomers, without requiring template assemblies or predefined composition. We benchmark StoPred against deep learning-based and template-based methods, and show that it achieves improved accuracy and efficiency across curated and blind datasets, with up to 16% and 41% higher top-1 accuracy for homomeric and heteromeric complexes, respectively, compared to the strongest prior method on our held-out test dataset. More importantly, StoPred is the first deep learning-based method capable of accurately predicting the stoichiometry of hetero-oligomeric complexes.

## 1 Introduction

Across all domains of life, proteins frequently function as part of larger assemblies composed of multiple polypeptide chains [1]. These protein assemblies represent an essential layer of cellular organization. They play indispensable roles in virtually every aspect of cell physiology, including enzymatic function [2], signal transduction [3], gene regulation [4], molecular transport [5], and structural scaffolding [3]. Assemblies can involve identical subunits (homo-oligomers) or different subunits (hetero-oligomers), and their formation can be essential for protein stability, folding [6] and function [1]. In many cases, the functional properties of a protein emerge only in the context of its quaternary structure [7], highlighting the centrality of oligomerization to biological function. Accordingly, deciphering the nature of protein assemblies is crucial for understanding the molecular basis of life. However, determining the assembly state and structure experimentally remains challenging and requires extensive effort[7, 8], which has become increasingly inadequate to meet the growing demands of the biomedical field. Fortunately, with the rapid advancement of artificial intelligence (AI) in recent years and its successful application in structural biology, the structures of most protein monomers and some protein assemblies can now be predicted with near-experimental resolution using powerful AI tools such as AlphaFold-Multimer [9, 10], AlphaFold3 [11], and DMFold [12]. This progress has, to some extent, alleviated the biomedical community’s high demand for experimental determination of high-resolution protein assembly structures. However, these AI tools require predefined stoichiometry to work properly, which poses a limitation when dealing with uncharacterized proteins. This challenge has attracted growing attention, particularly since CASP16 first introduced protein complexes with unknown stoichiometry as prediction targets[13].

Stoichiometry annotations in the Protein Data Bank (PDB) are currently based mainly on PISA algorithm predictions [14], with additional input from structure depositors. Although PISA predictions agree with experimental evidence on oligomeric states in approximately 90% of cases, the method relies on a solved crystal structure, which inherently limits its applicability [7, 15]. In the absence of experimental data, the inference of stoichiometry often relies on homology-based template searches against known assemblies, utilizing tools such as HMMER [16], MMseqs [17], and HH-suite [18]. However, the number of structurally characterized protein assemblies available in the PDB remains much smaller than the number of known protein sequences [19]. Therefore, the limited coverage of structurally characterized assemblies significantly restricts the applicability of template-based approaches for stoichiometry inference, especially for proteins without close homologs in the PDB. Protein multimer structure prediction methods, such as AlphaFold-Multimer and AlphaFold3, can be used to infer the stoichiometry of protein assemblies by inputting different stoichiometric combinations and evaluating the confidence scores of the resulting models to identify the most probable assembly composition (e.g., [20]). However, such approaches to stoichiometry prediction pose substantial computational challenges, as they require separate inferences for each possible copy number (which expands combinatorially with the number of distinct protein types present in the complex), and their reliability remains constrained by the quality of the multiple sequence alignment (MSA) or paired MSA [1]. MoLPC2 presents an approach that algorithmically determines the assembly mode of protein complexes based on AlphaFold-Multimer predictions [21]. This strategy is also computationally demanding, as it requires running all five parameterized models of AlphaFold-Multimer for each attempted stoichiometry combination [1].

Advances in natural language processing (NLP) have been successfully applied to protein sequences, giving rise to protein language models (pLM). These models are capable of learning structural and functional features embedded within amino acid sequences, including secondary structure, cell localization, and coevolutionary relationships [22–24]. Once trained, protein language models offer highly efficient inference, enabling their widespread application across numerous domains. In addition, Ora Schueler-Furman and colleagues demonstrated that such models are also able to capture information related to protein quaternary structure [25]. Therefore, more computationally efficient methods using pLMs for predicting stoichiometry have recently been developed, including QUEEN [7], DeepSub [26], and Syq2symm [1]. QUEEN is a sequence-based model that utilizes a pLM and a multilayer perceptron (MLP) to learn stoichiometric information embedded in ESM-2 representations [7]. DeepSub was subsequently proposed, leveraging both a protein language model and a bidirectional gated recurrent unit (Bi-GRU) to predict the stoichiometry of homo-oligomeric proteins [26]. Seq2Symm is a fine-tuned ESM-2 model that enables simultaneous prediction of both stoichiometry and symmetry for homo-oligomers [1]. However, these methods do not effectively consider template information even when it is available. More importantly, they are limited to predicting the stoichiometry of homo-oligomers, and are not applicable to hetero-oligomers, which constitute another major class of protein assemblies. These limitations constrain both the accuracy and the applicability of such methods.

Here, we present StoPred, a protein complex stoichiometry prediction method that combines protein language model embeddings with a graph attention (GAT) neural network [27] to enable subunit-level reasoning for both homo- and hetero-oligomeric complexes. By integrating both local and global predictions (defined in Methods 4.4), StoPred can infer stoichiometry directly from sequence or structure features, without requiring prior knowledge of the complex composition or template structures. We benchmark StoPred against existing sequence-based, template-based, and AlphaFold confidence score-based approaches, and show that it achieves higher accuracy and broader applicability on both curated and blind benchmark datasets, while also being much more computationally efficient. Our results indicate that StoPred fills a critical gap in computational modeling, providing an efficient tool for stoichiometry prediction of protein complexes with unknown or uncertain composition, and can also be used to guide the setup of high resolution modeling (e.g. using AlphaFold) in such cases.

## 2 Results

### 2.1 StoPred overview

StoPred is implemented as a deep learning model that uses protein language model embeddings as input and applies a graph attention network (GAT) to model inter-subunit relationships (Fig. 1). This GAT-based architecture constitutes the core StoPred framework, which we refer to as the StoPred GAT model in later sections. StoPred operates in four main steps. First, we represent each subunit type (i.e., each unique type of molecular entity) in a protein complex as a node embedding derived from protein language model features, capturing sequence or structure information. Then, we apply multiple layers of graph attention networks to propagate information between subunits, modeling inter-subunit relationships and updating each subunit’s representation in a permutation-invariant and equivariant manner. Finally, we use two prediction branches: a local branch that predicts the copy number distribution for each subunit type using a multilayer perceptron, and a global branch that predicts the overall stoichiometry state of the complex through a conditional mixture-of-experts module. The final stoichiometry prediction combines local and global outputs via a weighted scoring scheme to improve accuracy. Figure 1 illustrates the architecture, and the Methods section provides further details.

**Fig. 1.**
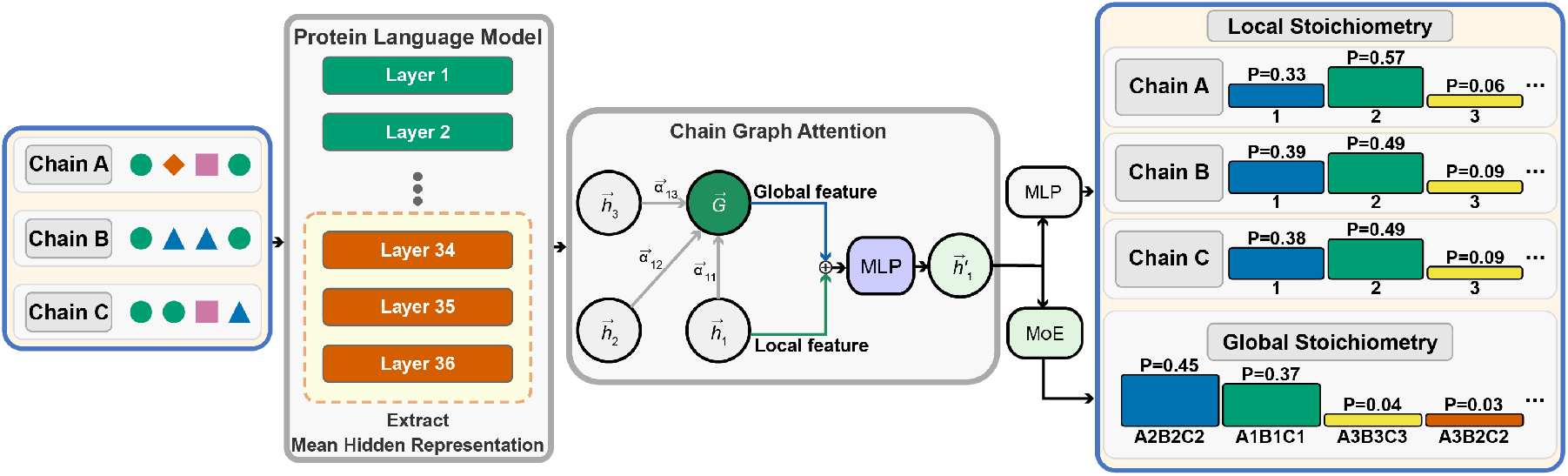
Overview of the StoPred GAT architecture. Protein sequences are first encoded into embeddings using a protein language model. These embeddings are then passed to a graph attention network, where each node represents a unique subunit type and edges represent potential inter-subunit interactions. Local and global branches predict subunit copy numbers and overall complex stoichiometry, and final predictions are obtained by combining outputs from both branches using a weighted scoring.

Throughout the discussion below, we use the notation “*AxBy*” (with *x, y* as integers) to denote a structure with two unique protein entity types, *A* and *B*, containing *x* copies of *A* and *y* copies of *B*. When referring to a specific monomer, *A*_*i*_ denotes the *i*th copy of protein entity *A*. The input to StoPred thus constitutes the identities (sequence, or optionally structure) of each unique entity type (*A, B*, etc.), and the output is a probability distribution of possible overall complex stoichiometries (*AxBy*).

### 2.2 PDB dataset evaluation

We benchmarked and compared our method against existing baseline methods using structures from the PDB database, divided by cutoff time (see Methods 4.1 for details). Since the distribution of the stoichiometry is imbalanced, we use several evaluation metrics, including accuracy, micro-F measure, and macro-F measure. For macro measurement, we aggregate all classes with fewer than 10 instances into a single “Other” class. The equations for micro-F and macro-F measures are provided in the supplemental file. Additionally, we report top-*N* accuracy (*i*.*e*., the fraction of targets for which the correct stoichiometry lies within the top *N* candidate stoichiometries, ranked by confidence score). In real applications, several stoichiometry candidates may be selected for downstream modeling, such as with AlphaFold3. Therefore, we also use top-*N* accuracy to measure the model’s ability to include the correct stoichiometry among the top-ranked candidates.

We trained two StoPred GAT models using either sequence (SP-sq-dl) or structure (SP-st-dl) features to examine the benefit of incorporating structural information in stoichiometry prediction. We also developed two alignment-based variants with the same features (SP-sq-al and SP-st-al) to serve as template-based baselines. All four models follow the naming format SP-{feature}-{method}, where “sq” or “st” indicates the input type (sequence or structure), and “dl” or “al” indicates the modeling approach (deep learning or alignment-based). Among these, we refer to the sequence-based deep learning model (SP-sq-dl) as StoPred, which we use as the default unless otherwise noted. We also compare our models with four existing or baseline methods: DeepSub, QUEEN, PreStoi, and “Naïve” (with the latter referring to predictions of each stoichiometry according to their existing frequencies in the training dataset). Note that DeepSub and QUEEN are designed for homomeric stoichiometry prediction, but are included here as baselines since they have been applied to heteromeric cases by independent subunit prediction [28]. Further details about each baseline are provided in the Methods section.

Figure 2 shows the evaluation results for all tested methods on the PDB dataset. The StoPred GAT models consistently outperform the baselines across accuracy, micro-F1, and macro-F1 (Fig. 2b), with the sequence-based and structure-based variants yielding nearly identical performance. In contrast, the StoPred alignment-based models perform worse than the GAT models but still substantially better than PreStoi, the other alignment-based method. All methods surpass the Naïve baseline. QUEEN and DeepSub, which are limited to homomeric stoichiometry prediction, achieve higher macro-F1 but perform worse than the Naïve baseline in terms of accuracy and micro-F1.

**Fig. 2.**
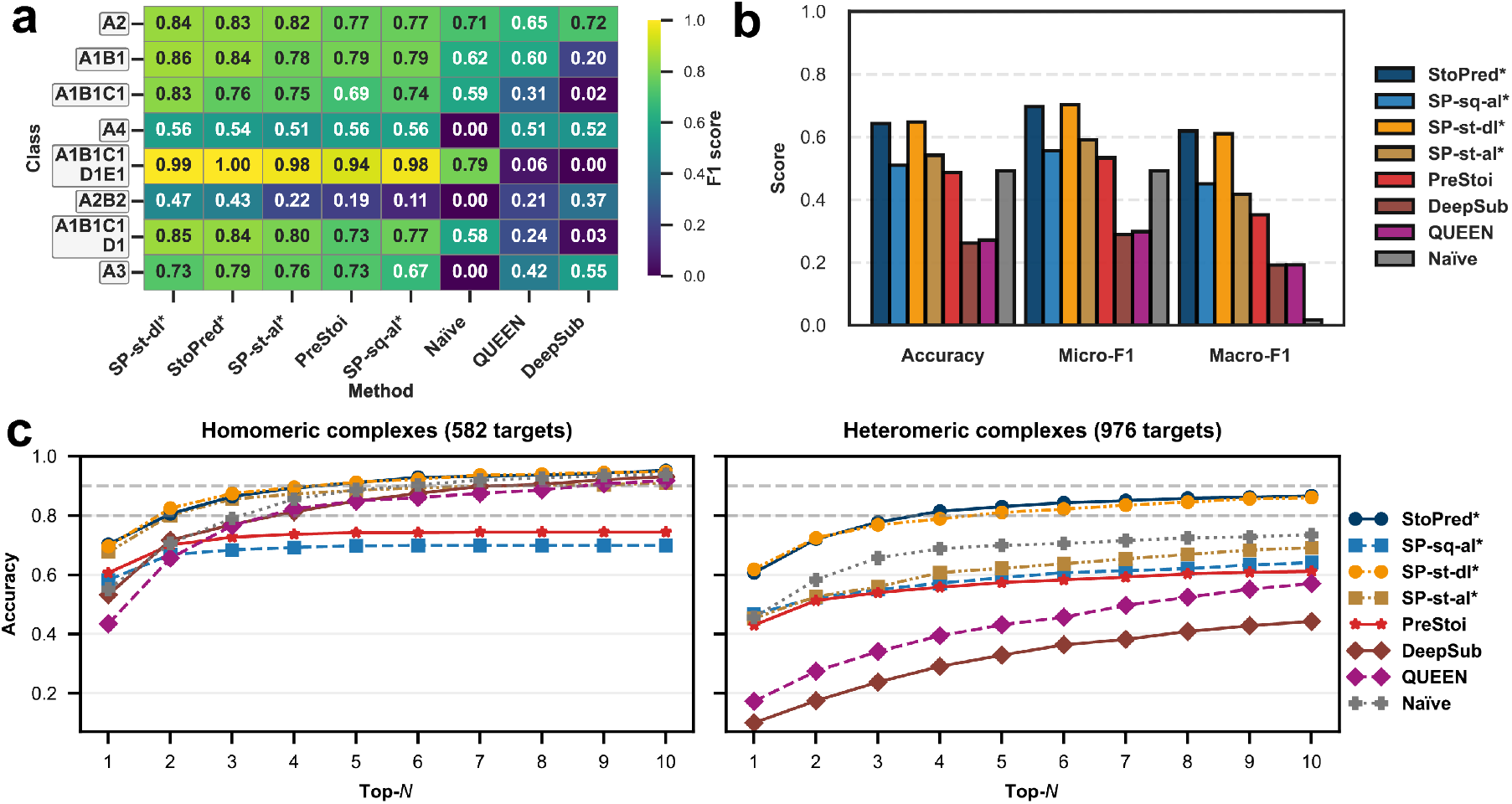
Performance comparison of StoPred models and baseline methods on the PDB dataset. (a) Heatmap showing prediction performance for the eight most frequent stoichiometry classes. (b) Overall accuracy, micro-F1, and macro-F1 for each method. (c) Top-*N* accuracy curves for all methods, evaluated separately on homomeric and heteromeric complexes. Methods marked with * are StoPred variants; “StoPred” here refers to the SP-sq-dl variant of our method.

The heatmap for the eight most frequent stoi-chiometry classes shows a similar trend (Fig. 2a), with the deep learning model providing a clear improvement over the alignment-based approach, especially for heteromeric classes such as A1B1 and A2B2. This improvement is likely due to the incorporation of the GAT mechanism, which enables the model to better capture interchain relationships. The top-*N* accuracy curves further illustrate the advantage of the StoPred GAT models (Fig. 2c), as they include the correct stoichiometry among the top predictions more often than the baseline methods. For homomeric complexes, the structure alignment model (SP-st-al) shows a modest improvement over the sequence alignment model (SP-sq-al), bringing its performance closer to the GAT models. However, this benefit does not extend to heteromeric complexes, likely because the current scoring function matches features at the chain level rather than considering the global similarity of the entire complex. In addition, the GAT models (SP-st-dl and SP-sq-dl) still uniformly outperform the corresponding alignment based methods. In contrast, within the StoPred GAT models, there is no such trend between sequence and structure inputs, likely because both types of features are derived from ESM-family models and thus provide similar representations.

For heteromeric complexes, as *N* increases, the Naïve method improves rapidly and eventually outperforms all other baseline methods. However, the StoPred GAT models consistently provide the highest quality top-*N* candidate predictions across all tested *N*.

Considering both computational cost and predictive performance, we use the sequence-based deep learning model as the final StoPred implementation. For subsequent comparisons, we focus on the sequence-feature variants (SP-sq-dl and SP-sq-al) and do not include the structure-based models.

To compare the performance of StoPred with AlphaFold3, we used a filtered subset of the PDB time-based test set due to computational constraints. This subset contained 255 homomeric and 642 heteromeric complexes (detailed in Sec. 4.1). Only sequence features were used for StoPred in this comparison, as we selected sequence-based features for our final model. Other methods are included for reference. We also restricted AlphaFold3 to consideration of no more than six stoichiometries: the five top stoichiometries from StoPred, plus the correct stoichiometry (if not already included in the StoPred top-5), thus limiting the computational effort required and providing a biased evaluation in favor of AlphaFold3.

Figure 3 summarizes the comparison between StoPred and AlphaFold3 on the selected dataset. The StoPred GAT model achieves the highest performance across accuracy, micro-F1, and macro-F1 (Fig. 3b), followed by the AlphaFold3 ranking score model, which is slightly better than the StoPred alignment-based model. Both perform better than PreStoi and the other baseline methods. The F1 score heatmap for the eight most frequent stoichiometry classes (Fig. 3a) shows that both StoPred and AlphaFold3 achieve the highest F1 scores for most classes, with each model showing strengths in different stoichiometry types. The top-*N* accuracy curves are shown in Fig. 3c. As noted above, our test procedure for AlphaFold3 guarantees 100% accuracy at top-6 and thus gives it an advantage at higher N values. Despite this bias, StoPred outperforms AlphaFold3 at top-1 and top-2, and achieves comparable accuracy at top-3 for both homomeric and heteromeric complexes. Additionally, StoPred not only achieves higher prediction accuracy than AlphaFold3, but also requires orders of magnitude less computational time (Fig. 3d). This panel shows the runtime for generating the final prediction with StoPred compared to complex modeling with AlphaFold3. Notably, for AlphaFold3 stoichiometry prediction, multiple candidate models must be generated, and larger residue counts further increase runtime. Together, these results show that StoPred GAT provides higher prediction accuracy and greater computational efficiency than AlphaFold3 for stoichiometry prediction on our restricted PDB benchmark.

**Fig. 3.**
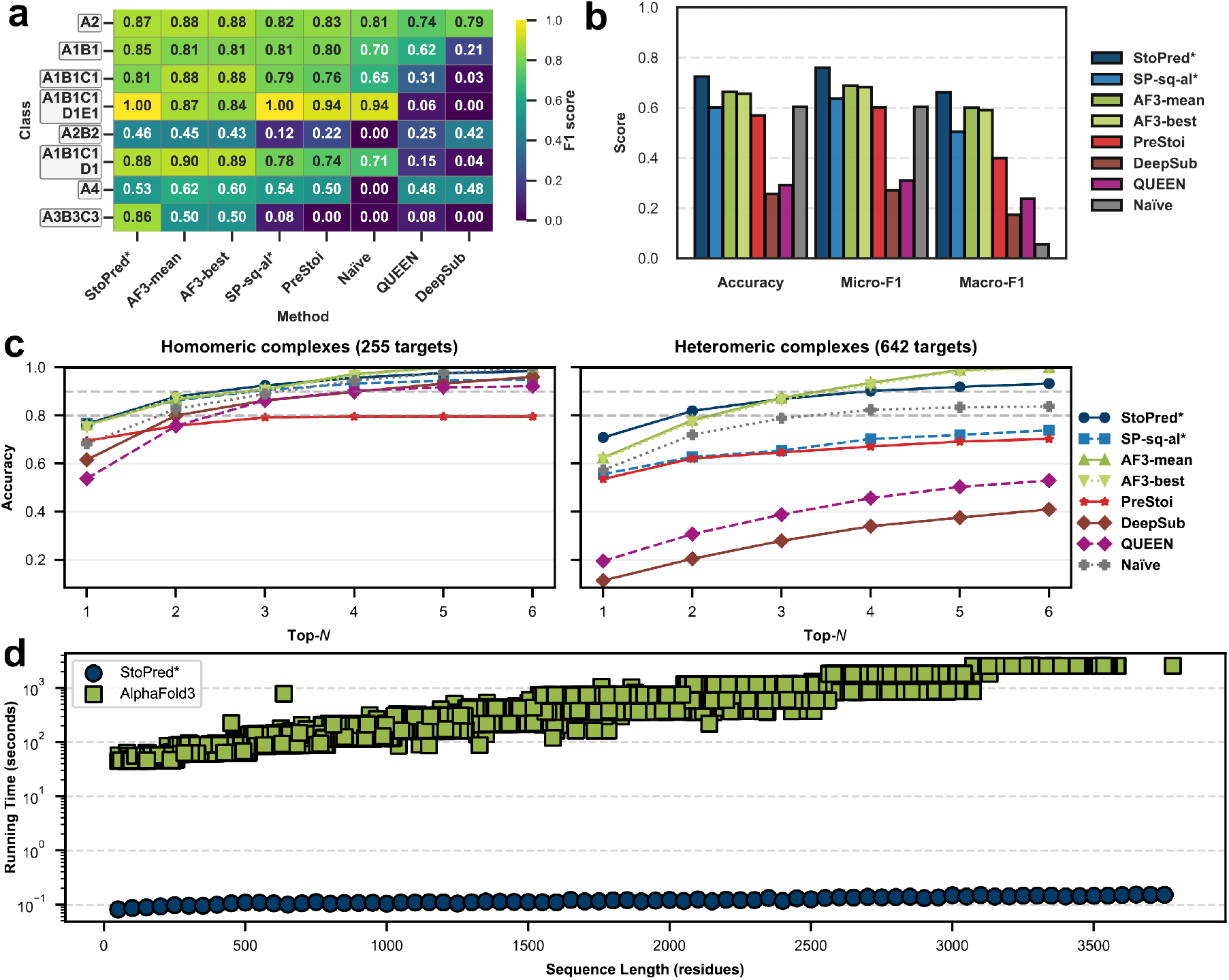
Performance comparison of StoPred models, AlphaFold3, and baseline methods on the AF3-PDB dataset. (a) Heatmap showing prediction performance for the eight most frequent stoichiometry classes. (b) Overall accuracy, micro-F1, and macro-F1 for each method. (c) Top-*N* accuracy curves for all methods, evaluated separately on homomeric and heteromeric complexes. (d) Runtime comparison for final stoichiometry prediction by StoPred and structure modeling by AlphaFold3. Methods marked with * are StoPred variants.

### 2.3 CASP16 Phase 0 dataset evaluation

To further evaluate the performance of our method, the baseline methods, and the current leading human predictors in real-world challenges, we used data from the “phase 0” evaluation of the recent CASP16 experiment. In this benchmark, the targets are protein structures that were newly determined but not yet publicly released. Since both our training and validation datasets only include proteins released before the start of CASP16, this benchmark represents a challenging and unbiased evaluation for stoichiometry prediction.

Figure 4 shows the evaluation results. As shown in Fig. 4b, we found that the SP-sq-al method, which relies heavily on sequence similarity, performed much worse compared to its performance on the PDB dataset. Although PreStoi also relies on sequence information, it performed slightly better than SP-sq-al on this dataset, likely because multiple sequence alignment profiles are more sensitive for detecting distinct templates. AlphaFold3 provides a slight improvement over these methods, but because most CASP targets are very large and exceeded the memory capacity of our benchmarking hardware (48GB), AlphaFold3 fails to model some stoichiometry candidates at all, resulting in lower overall performance. By contrast, StoPred maintains strong performance and provides a larger performance gap compared to other methods. The F1 score heatmap for the most frequent classes (Fig. 4a) shows that StoPred (SP-sq-dl) achieves the best F1 score for all classes except for A4. In terms of top-*N* accuracy (Fig. 4c), StoPred again achieves the best results. Note that for the CASP16 dataset, due to the small number of targets, we did not group classes with fewer than 10 instances into an “Other” class when calculating micro-F1 and macro-F1 scores in Fig. 4b.

**Fig. 4.**
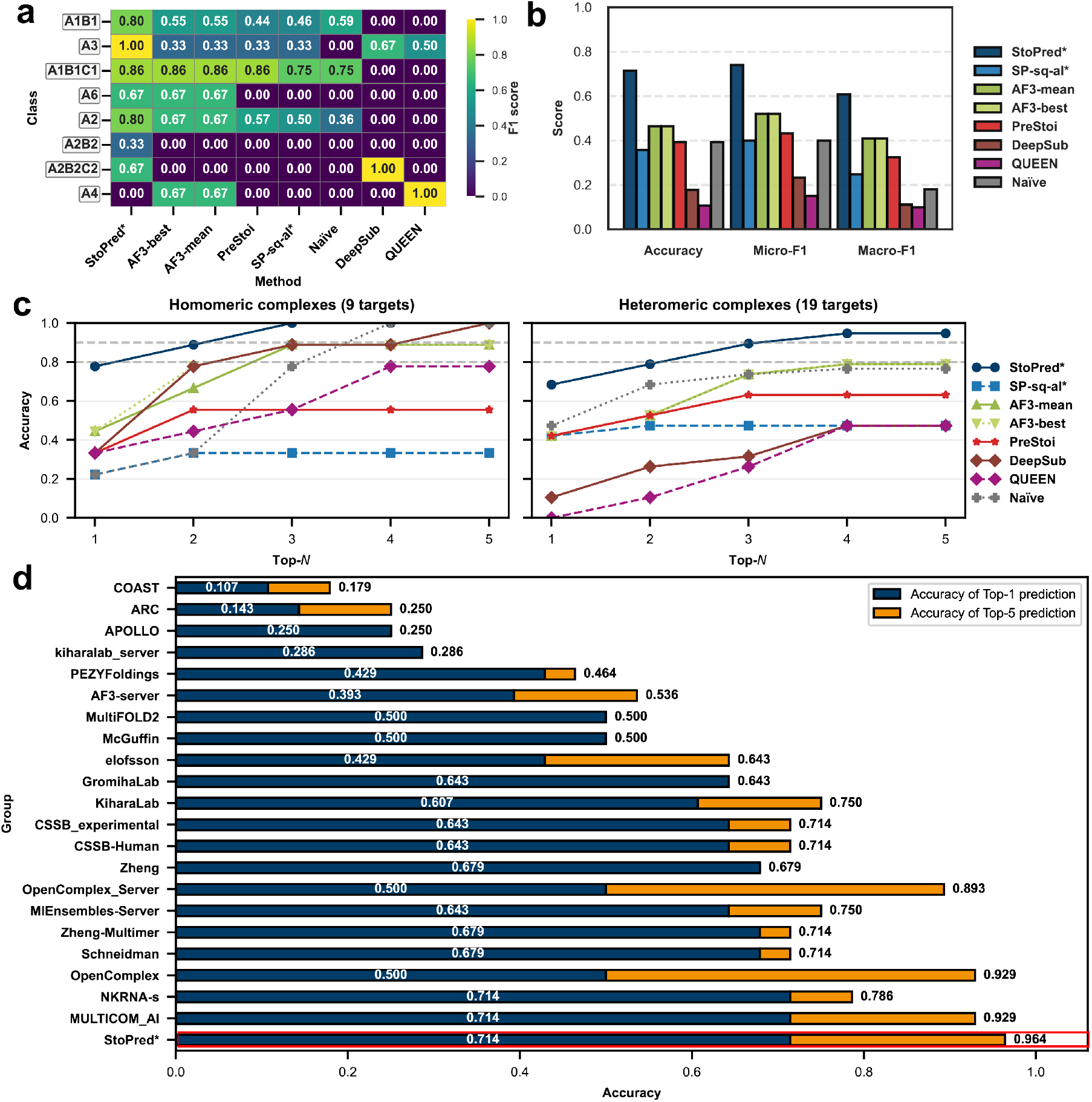
Performance comparison of StoPred models, AlphaFold3, and other baseline methods on the CASP16 phase 0 dataset. (a) Heatmap showing prediction performance for the eight most frequent stoichiometry classes. (b) Overall accuracy, micro-F1, and macro-F1 for each method. (c) Top-*N* accuracy curves for all methods, evaluated separately on homomeric and heteromeric complexes. (d) Comparison of top-1 and top-5 accuracy on the CASP16 benchmark, including StoPred (SP-sq-dl; highlighted in red) and CASP16 participants (Data from the official CASP16 website). Throughout the figure, methods marked with * are StoPred variants.

When compared with CASP16 participants (Fig. 4d), StoPred (SP-sq-dl) ties with NKRNA-s and MULTICOM AI for top-1 accuracy at 0.714. As predictors in CASP are permitted to submit up to five models for each target, we also report top-5 accuracy in this evaluation, where SP-sq-dl outperforms all predictors with an accuracy of 0.964. These findings suggest that StoPred learns stoichiometry information from sequence features effectively and allows accurate stoichiometry prediction even for challenging targets.

### 2.4 Ablation Study

To evaluate the contribution of each component in our model, we performed an ablation study by removing specific modules and measuring the resulting effects on the final model performance. Here, StoPred represents our complete model, and the variants with “noGAT” or “noGlob” indicate removal of the corresponding component. Specifically, “noGAT” refers to disabling the graph attention network for modeling inter-chain relationships, while “noGlob” refers to excluding the global prediction module. We only applied the global prediction module in heteromeric complex prediction, as for homomeric complexes the global prediction is equivalent to the subunit-level prediction since there is only a single unique subunit. The model lacking both the global and GAT components (i.e., “noGAT-noGlob”) is thus very similar in spirit to existing methods such as QUEEN, and the ablation study permits us to assess the contributions of the new components added in StoPred.

Figure 5 summarizes the results. The overall performance differences in the ablation study are more pronounced for heteromeric complexes than for homomeric complexes (Fig. 5c). For homomeric complexes, the GAT feature clearly drives improved accuracy, micro-F1, and macroF1, likely because the graph attention network helps the model distinguish between identical chains and reduces confusion. For heteromeric complexes, the global prediction module becomes increasingly important as the number of candidate states considered (top-*N*) increases. Removing the GAT has the largest impact on top-1 accuracy for heteromers, but as N increases, the global prediction module has the largest effect on top-*N* accuracy (Fig. 5a); importantly, however, we observe that both the GAT and global components make important contributions to the predictions, as the performance of the noGAT-noGlob model is substantially lower than that of models lacking either of those two components independently.

**Fig. 5.**
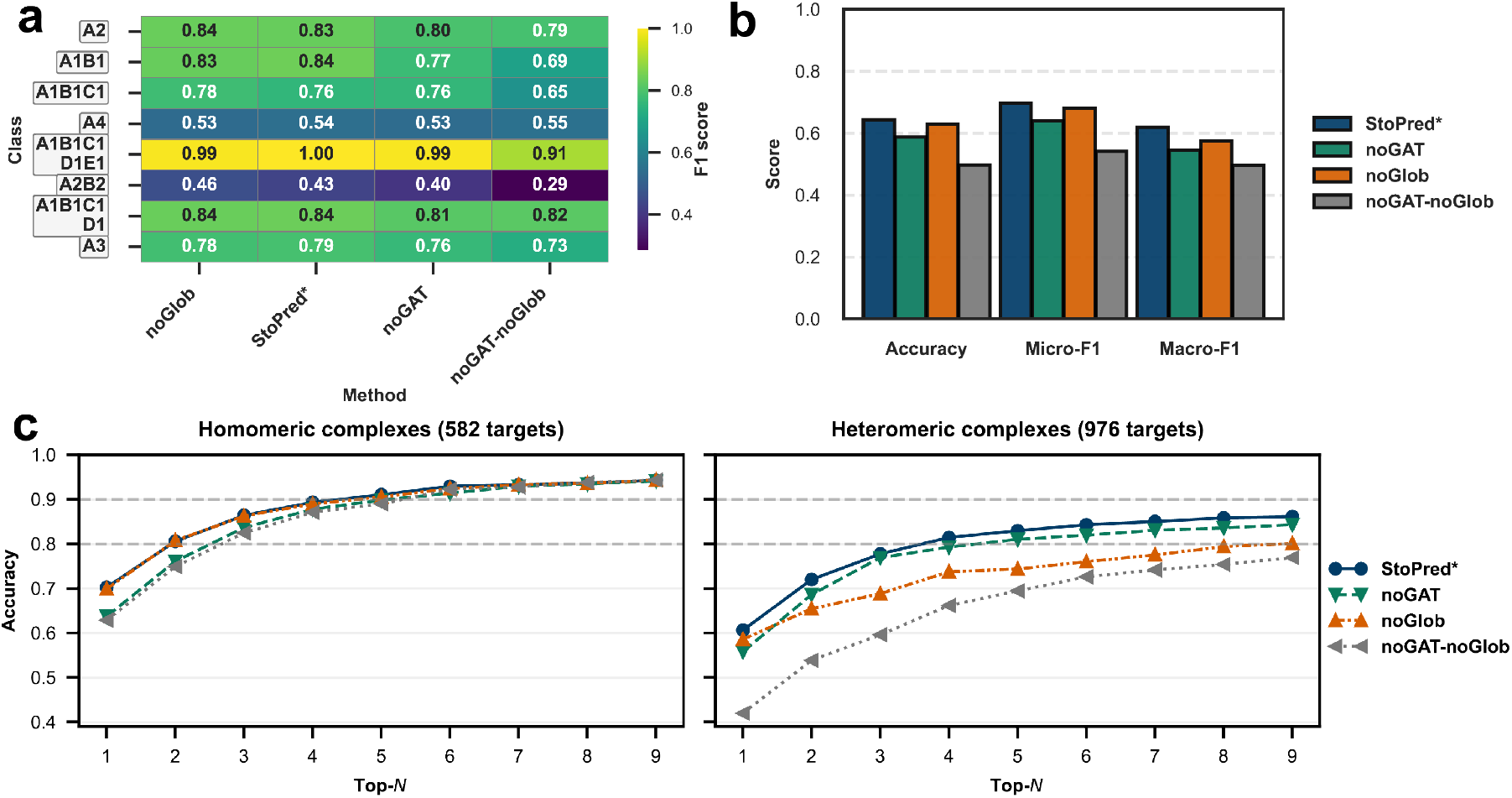
Ablation study evaluating the contribution of the graph attention network and global prediction module in StoPred on the PDB dataset. (a) Heatmap showing prediction performance for the eight most frequent stoichiometry classes. (b) Overall accuracy, micro-F1, and macro-F1 for each model variant. (c) Top-*N* accuracy curves for all models, evaluated separately on homomeric and heteromeric complexes.

Overall, the ablation study shows that both the GAT and the global prediction module are essential for optimal performance. The GAT improves accuracy by modeling inter-chain relationships, while the global prediction module is especially important for identifying the correct stoichiometry among the top-*N* heteromeric complex candidates. The contributions of both major modules of StoPred to heteromeric stoichiometry prediction appear especially important given the higher complexity and biological importance of heteromeric interactions, and the relative lack of alternative tools for this prediction task.

### 2.5 Case studies on the advantage of StoPred over AlphaFold3 score-based selection

To further investigate the advantages of the StoPred GAT model over the commonly used AlphaFold3 ranking score-based selection approach, we conducted a case study on two representative protein complexes from our test dataset: *Staphylococcus aureus* MazF in complex with nanobody 4 (PDB ID: 9G2A) and *Lactococcus lactis subsp. lactis* CdaA-DAC domain in complex with GlmM (PDB ID: 9G69) [29]. Both complexes are heterotetramers with a stoichiometry of A2B2. Figures 6 and 7 summarize the results, including structure alignments between predicted models and experimentally determined structures, confidence score heatmaps for StoPred and AlphaFold3, and predicted aligned error (PAE) heatmaps for both the highest ranking AlphaFold3 model and the AlphaFold3 model with the correct stoichiometry.

**Fig. 6.**
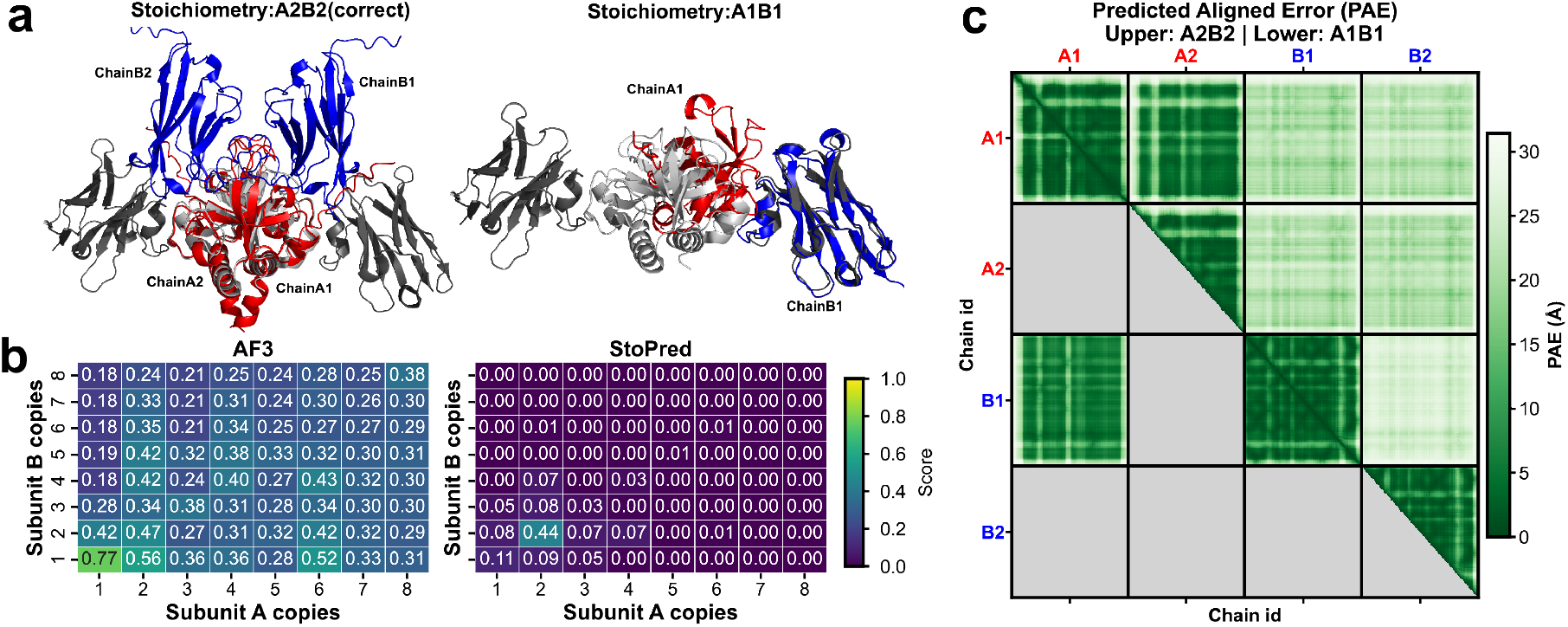
Case study comparing StoPred and AlphaFold3 stoichiometry prediction for the *Staphylococcus aureus* MazF–nanobody 4 complex (PDB ID: 9G2A). (a) Structural alignment of predicted models to the native structure: predicted A, blue; predicted B, red; native A, light grey; native B, dark grey. The A2B2 model (correct stoichiometry) and the A1B1 model (highest AlphaFold3 ranking score) are shown. (b) Heatmaps showing ranking scores for different stoichiometry candidates with AlphaFold3 and confidence scores predicted by StoPred. (c) Predicted aligned error heatmap for AlphaFold3 models: the upper right triangle shows the PAE for the true A2B2 stoichiometry, and the lower left triangle shows the PAE for the highest scoring A1B1 model.

**Fig. 7.**
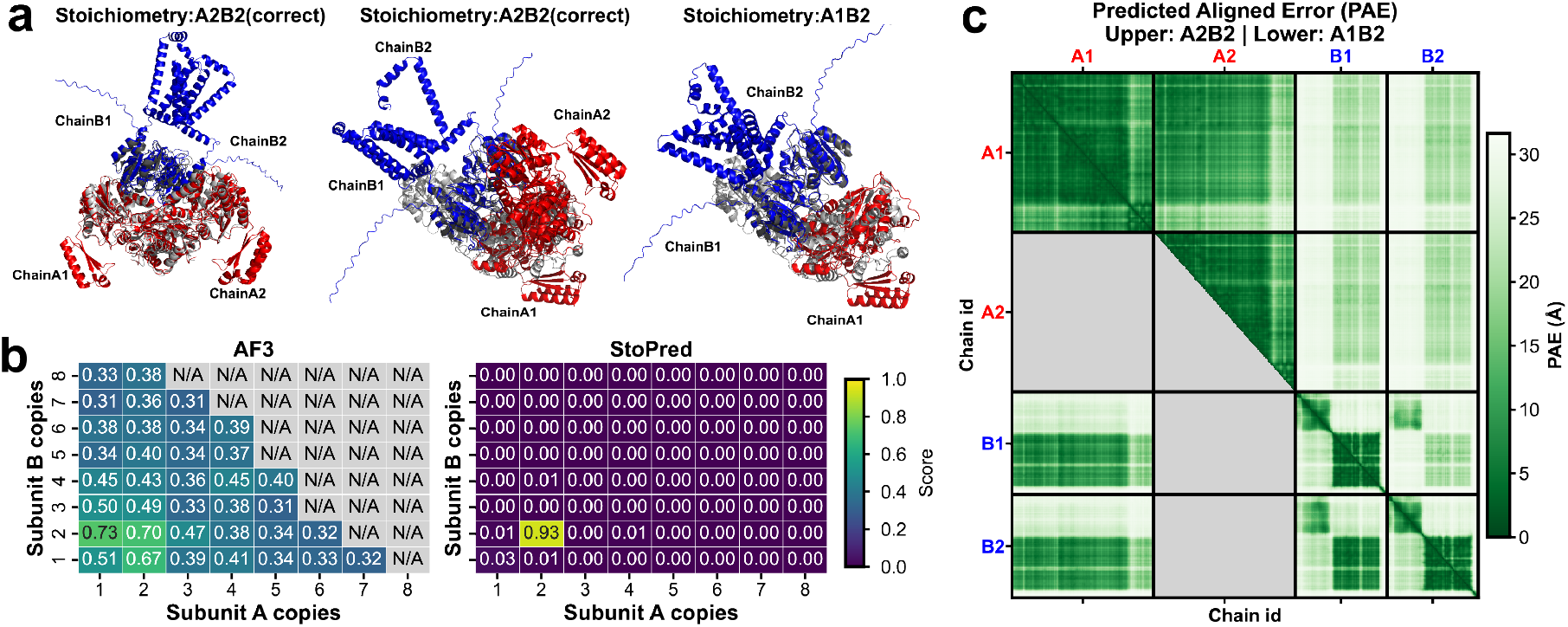
Case study comparing StoPred and AlphaFold3 stoichiometry prediction for the *Lactococcus lactis subsp. lactis* CdaA-DAC–GlmM complex (PDB ID: 9G69). (a) Structural alignments of predicted models to the native structure: left panel shows the A2B2 model aligned by native *A*_1_*A*_2_ chains, middle panel shows the A2B2 model aligned by native *B*_1_*B*_2_ chains, and right panel shows the A1B2 model. Predicted A: blue, predicted B: red, native A: light grey, native B: dark grey. (b) Heatmaps of ranking scores for different stoichiometry candidates using AlphaFold3 (with N/A values for candidates where models could not be generated due to VRAM limitations) and confidence scores from StoPred. (c) Predicted aligned error heatmap for AlphaFold3 models, with the upper right triangle showing the PAE for the true A2B2 stoichiometry and the lower left triangle showing the PAE for the highest scoring alternative model.

For 9G2A, AlphaFold3 assigns the highest ranking score (0.77) to a model with A1B1 stoichiometry, while the true A2B2 stoichiometry receives a lower score (0.56) (Fig. 6b). Structural alignment of the A2B2 model to the native structure reveals that AlphaFold3 can correctly model the interaction between chain *A*_1_ and chain *A*_2_, but fails to position chains *B*_1_ and *B*_2_ accurately (Fig. 6a), as reflected in the PAE heatmap (Fig. 6c). This demonstrates that the ranking score primarily reflects the model quality for the generated structure, rather than the correctness of the underlying stoichiometry. In contrast, StoPred, through integration of global and local predictions, correctly identifies the true A2B2 stoichiometry as the top candidate, with a confidence score of 0.44.

For 9G69, StoPred again predicts the correct A2B2 stoichiometry with a high confidence score (0.93), whereas AlphaFold3 yields similar ranking scores for A1B2, A2B1, and A2B2 models (Fig. 7b), making it difficult to identify the correct stoichiometry based on ranking score alone. Structural alignment indicates that the input sequence includes an extended C-terminus for subunit B, which is absent in the solved structure (Fig. 7a). This may cause AlphaFold3 to misestimate the PAE between chain *A*_1_ and chain *B*_1_ (Fig. 7c). Nevertheless, alignment of chain pairs *A*_1_*A*_2_ and *B*_1_*B*_2_ to the corresponding native subunits is accurate, but the relative arrangement between the two pairs is slightly offset. This further illustrates that relying on the AlphaFold3 ranking score alone is not optimal for determining stoichiometry.

## 3 Discussion

StoPred is a protein complex stoichiometry prediction method that improves prediction performance for both homomeric and heteromeric complexes. StoPred addresses the gap in heteromeric complex stoichiometry prediction by combining protein sequence features from pretrained protein language models, message passing between subunit embeddings using a graph attention network, and a decomposition approach that considers both local subunit-level and complex-level information. Our results suggest that protein language model embeddings capture quaternary structure information and support direct classification of complex stoichiometry. The graph attention network allows StoPred to model dependencies between subunits in heteromeric complexes, which are missed by models treating subunits independently. Decomposing the prediction into local and global components helps balance subunit-specific and complex-level information, leading to improved accuracy and faster runtime for heteromeric stoichiometry prediction. Although current structural modeling pipelines typically use AlphaFold to generate and rank multiple candidate stoichiometries, the AlphaFold confidence metrics (such as ipTM or pTM) are only shown to correlate with structural accuracy and not with the correct stoichiometry itself. StoPred sidesteps this limitation by predicting copy numbers directly from sequence or structure features, without requiring expensive structure generation. As a result, StoPred can quickly suggest a small set of likely stoichiometry candidates, which can be used to narrow down structural modeling with AlphaFold or similar tools. While StoPred already provides improved efficiency and accuracy over current approaches, there are areas for future improvement. We aim to design improved structure-based embeddings to capture features not available in current representations, and with these better input encodings, to develop multi-modal models that integrate template information, sequence, and structural features in a unified framework. At present, StoPred is limited to protein-only complexes and assumes that each input chain type is present at least once in the biologically correct assembly; it is therefore not suitable for predicting the presence versus absence of interactions between different molecular entities. Extending the method to protein–nucleotide complexes will be a focus of future work, as will extensions to provide information about the absence of interactions.

Overall, StoPred is able to predict stoichiometry directly and efficiently, and when used together with structure modeling tools, it allows for fast modeling of large protein complexes from sequence alone. This makes StoPred a practical tool for complexes with unknown composition and supports downstream structural and functional analysis.

## 4 Methods

### 4.1 PDB dataset

To train and evaluate our model for protein complex stoichiometry prediction, we constructed a large-scale dataset from the Protein Data Bank (PDB) [30]. We downloaded all non-obsolete mmCIF files from the PDB archive and extracted biological assemblies as defined by the original authors. Only assemblies composed entirely of ‘polypeptide(L)’ chains were retained, and for assemblies with identical subunit composition and stoichiometry, only one representative was kept to reduce redundancy. This resulted in a dataset of 63,142 unique author-defined biological assemblies.

To reflect how new protein complexes are encountered in practice, we adopted a time-based split for training, validation, and testing, similar to the strategy used in the CAFA challenge [31]. Assemblies deposited before October 1, 2023, were used for training. Those deposited between October 2, 2023, and May 1, 2024, formed the validation set. For the test set, we excluded biological assemblies that lacked experimental support to ensure high data quality. To remove homology contamination, we applied sequence-based redundancy filtering using MMseqs [32]. For each test complex, chains were optimally mapped to those in the training and validation sets. A test complex was considered redundant and removed if all of its chains could be mapped with sequence identity greater than 0.8 to chains in a training or validation complex. Complexes with at least one unmapped or low-identity chain were retained. After this filtering, the test set contained 582 homomeric complexes and 976 heteromeric complexes (reduced from 1,251 and 1,457, respectively).

For benchmarking StoPred against AlphaFold3, we applied additional filtering to the test set based on complex size and subunit copy number to fit computational constraints. Specifically, we selected complexes with a total amino acid length less than 4,200, a maximum singlechain length less than 500, and no more than 8 copies per subunit. After applying these criteria, the filtered test set contained 255 homomeric complexes and 642 heteromeric complexes. Detailed stoichiometry class distributions for all datasets are provided in the supplementary information.

### 4.2 CASP16 dataset

To fairly assess the performance of our model in realistic settings and in direct comparison with current leading approaches, we used the Phase 0 dataset from the sixteenth Critical Assessment of protein Structure Prediction (CASP16) experiment [13]. CASP is a well-established community experiment designed to benchmark protein structure and complex prediction methods.

During Phase 0, 28 multimeric protein targets were released to participants without any stoichiometry information. These included 9 homomultimers and 19 hetero-multimers. For each target, predictors were required to submit both the predicted stoichiometry and structural models generated based on their predicted stoichiometry. This setup closely resembles real-world scenarios, where the composition of a protein complex is not always known in advance.

For our evaluation, we used all 28 Phase 0 targets and the stoichiometry predictions submitted by the expert groups. These targets span a variety of oligomerization states, including both well-characterized and more challenging complexes, making this dataset a strong albeit small test for stoichiometry prediction models.

### 4.3 Protein language model ESM

We leveraged state-of-the-art Evolutionary Scale Model (ESM)[33] family protein language models to derive informative representations for each subunit. ESM models are transformer-based language models trained on large protein sequence databases, and have been shown to produce sequence representations that are predictive for a range of biochemical and structural properties of proteins.

We use esmc-600m-2024-12 [34] to generate protein sequence representations for all proteins in our dataset. Specifically, we compute the hidden states from the last three layers of the model. For each layer, the per-residue embeddings are averaged across the amino acid sequence, resulting in a final embedding of size 3 × 1,152 for each protein subunit.

For structure-based features, we first generate a predicted 3D structure for each subunit using AlphaFold3. The predicted atomic coordinates are then processed by the esm3-sm-open-v1 [35] model, which contains 1.4 billion parameters, to extract a structure-informed embedding for each subunit. For these features, we compute the hidden state from the last layer of the model and average the per-residue embeddings across the sequence, resulting in an embedding of size 1,536 for each protein subunit.

### 4.4 StoPred deep learning model

In StoPred GAT model (deep learning-based, or “dl”), we use ESM familiy models to represent protein feature and with a graph attention architecture for subunit-level reasoning. Each protein complex is modeled as a set of subunits, where each subunit is a node and edges represent potential inter-subunit relationships. Each node is initialized with a fixed-dimensional embedding derived from either sequence or structure features.

Given a protein complex with *C* unique subunits, let **X** *∈* ℝ^*C×d*^ denote the matrix of input subunit embeddings, where *d* is the embedding dimension. Each node embedding **x**_*c*_ is first transformed by a multi-layer perceptron (MLP):

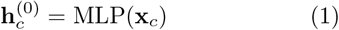

The core of the model is a multi-head graph attention layer. For each attention head *h*, node features are projected as 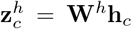. Additive attention coefficients are computed for each pair of subunits:

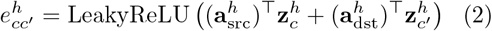

Here, LeakyReLU is a nonlinear activation function, 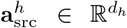and 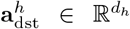are learnable parameters for each attention head, where *d*_*h*_ is the dimension per head. These parameters are used to independently transform the source and target node representations before calculating their pairwise attention scores. Self-attention terms and positions corresponding to padded subunits are masked out so that only valid inter-subunit pairs are considered. Attention weights are then normalized over all valid neighbors through a softmax operation.

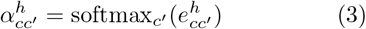

The aggregated feature for node *c* is then given by

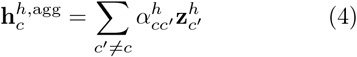

The outputs of all heads are concatenated, followed by a linear layer, ELU activation, dropout, and a residual connection:

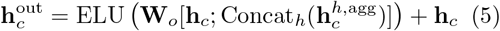

This operation is repeated for multiple layers. Masking is applied throughout to handle variable complex sizes.

To capture stoichiometry at different levels, we divide prediction into two branches: a local branch, which assigns a copy number to each individual subunit type, and a global branch, which predicts the overall composition of the complex as the composition of copy numbers across all subunits (e.g., (3, 2) corresponds to either *A*2*B*3 or *B*3*A*2).

For global stoichiometry prediction, we use a conditional mixture-of-experts (MoE) module. Experts are selected according to the number of valid subunits in the complex, and each expert predicts the logits for the corresponding set of possible global stoichiometries. In addition, a shared expert is included to handle cases where the subunit number does not match any predefined configuration, as well as to predict an “other” class for global stoichiometry states that are rare or not present in the training data:

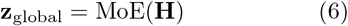

where **H** is the set of final subunit representations for the complex.

For local copy number prediction, each subunit’s representation is passed to a classification layer:

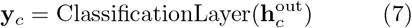

A softmax is applied to produce the predicted distribution over copy numbers for each chain.

For heteromeric complexes, we combine both local and global predictions to improve the accuracy of top-*K* stoichiometry assignments. For each candidate stoichiometry state, the local probability is computed by multiplying the predicted probabilities for each subunit copy number across all chains. The global probability for the same state is directly obtained from the output of the MoE module. The final score for each candidate state is calculated as a weighted sum:

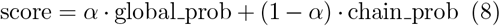

where *α* is a weighting parameter reflecting the importance of the global prediction relative to the product of local probabilities. The value of *α* is selected based on performance on the validation dataset. Details of the *α* selection and evaluation are provided in the Supplementary Information.

The model is trained with cross-entropy loss for both the local and global heads. Further architectural details, including regularization and optimization procedures, are provided with the code release.

### 4.5 StoPred template-based prediction

In StoPred template-based prediction (alignment-based, or “al”), we use a template database constructed from the training dataset to infer stoichiometry for new protein complexes. The pipeline is implemented for both sequence-based and structure-based templates, sharing similar overall steps but differing in the feature type and matching strategy.

For the sequence-based template approach, we extract subunit sequences from the mmCIF files in the training set and construct a sequence template database using MMseqs2[32]. For the structure-based template pipeline, we generate predicted structures for all training subunits using AlphaFold3, and use Foldseek[36] to build a structure template database from these models.

The underlying hypothesis is that protein complexes with similar subunit composition tend to share similar stoichiometry states. For a given query complex with *C* subunits, we search each subunit against the corresponding template database using MMseqs2 for sequences or Foldseek for structures, and collect the reported *K* hits for each chain. For each candidate copy number state *y*, we compute a confidence score from the *K* hits as follows:

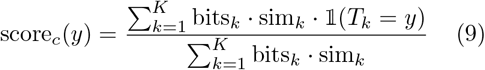

where bits_*k*_ is the bitscore of the *k*-th template hit to the query chain *c*, sim_*k*_ is the similarity between the query chain and the template chain, measured by sequence identity for the sequence-based pipeline and TM-score for the structure-based pipeline. When calculating sequence identity or TM-score, normalization is based on the larger chain length if the query and template differ. 𝟙(*T*_*k*_ = *y*) is an indicator function equal to 1 if the template *T*_*k*_ has *y* copies of the subunit, and 0 otherwise.

### 4.6 Naïve appoarch

Because stoichiometry class distribution is highly imbalanced, it is possible to obtain prediction results by simply assigning stoichiometry states according to their observed frequencies within the training set. For a given number of subunits, we compute the frequency of each stoichiometry state *S* as

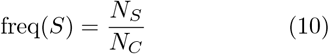

where *N*_*S*_ is the number of training protein complexes that have both stoichiometry class *S* and subunit count *C*, and *N*_*C*_ is the total number of training protein complexes with subunit count *C*.

For evaluation, we rank stoichiometry states by their observed frequencies in the training set for each subunit count and select the top-ranked states as predictions. When the predicted state is not a homomer, the subunit order is randomly shuffled to avoid bias, and the results are averaged across 1000 runs. If the test complex has a subunit count not present in the training data, we predict a state with one copy of each chain.

### 4.7 PreStoi

PreStoi[20] is a template-based method for stoichiometry prediction that searches for homologous complexes in the Protein Data Bank using sequence alignments. For each complex, PreStoi outputs a list of candidate copy numbers for each unique subunit, based on the occurrence of similar subunits in homologous templates. If sufficient template evidence is found, PreStoi also provides a final template-based prediction as its recommended stoichiometry. To benchmark PreStoi, we assigned per-chain confidence scores using the empirical frequency of each copy number among the template suggestions. When a final template-based prediction was available, we assigned it the highest confidence. When no template information was available, we provided a prediction with one copy of each chain.with one copy of each chain.

### 4.8 AlphaFold3 score-based selection

AlphaFold3 outputs a ranking score for each predicted structure, capturing both the overall model quality and the confidence in predicted subunit interfaces. To infer the most likely stoichiometry for a protein complex, we enumerate possible subunit copy number combinations and generate structural models for each such candidate stoichiometry. For each candidate, we use either the maximum (AF-best) or the mean (AF-mean) AlphaFold3 ranking score from the 5 generated models as a confidence estimate for that stoichiometry.

Due to the high computational cost of AlphaFold3, our benchmark is limited to a maximum of six stoichiometry candidates per complex. Specifically, we include the top five predictions from the StoPred sequence deep learning model, along with the true stoichiometry if it is not already among the top five. While this approach is designed to facilitate a direct comparison, it introduces a bias in favor of AlphaFold3 over StoPred, as including the most probable candidates and the true stoichiometry matches or exceeds the performance expected if a larger set of candidates were evaluated.

### 4.9 DeepSub

DeepSub [26] is a sequence-based method for predicting protein oligomerization states. It uses ESM2 protein language model embeddings, a bidirectional gated reccurent unit (Bi-GRU), and an attention layer to predict the stoichiometry of homo-oligomers from protein sequences alone. Since there are currently no deep learning models available for hetero-oligomer stoichiometry prediction, we include DeepSub as a comparison by applying it to each subunit sequence in a given complex. For hetero-oligomer benchmarking, we aggregate the predicted probabilities for each chain by using beam search to generate the top-*N* candidate stoichiometries, ranking them according to the product of the chain-level probabilities. This allows for direct comparison with our method.

### 4.10 QUEEN

QUEEN[7] is a sequence-based method similar to DeepSub, but replaces the Bi-GRU and attention layer with an MLP. We use the same aggregation strategy as described for DeepSub to extend QUEEN to hetero-oligomer stoichiometry prediction.

## 5 Data availability

All data underlying this work is freely available at https://seq2fun.dcmb.med.umich.edu/StoPred/StoPred.tar.xz

## 6 Code availability

The source code of this work is freely available at https://github.com/QuanEvans/StoPred

## Supplementary information

If your article has accompanying supplementary file/s please state so here.

Authors reporting data from electrophoretic gels and blots should supply the full unprocessed scans for key as part of their Supplementary information. This may be requested by the editorial team/s if it is missing.

Please refer to Journal-level guidance for any specific requirements.

## Acknowledgements

This work was supported by the Advanced Cyberinfrastructure Coordination Ecosystem: Services & Support (ACCESS) program, funded by the National Science Foundation (2138259, 2138286, 2138307, 2137603, and 2138296); the National Institute of Allergy and Infectious Diseases (AI134678 to L.F.); the National Natural Science Foundation of China (12426303 to W.Z.); the Fundamental Research Funds for the Central Universities (054-63253109 to W.Z.); and the Tianjin Science and Technology Program (24ZXZSSS00320 to W.Z.).

